# Neurobiological successor features for spatial navigation

**DOI:** 10.1101/789412

**Authors:** William de Cothi, Caswell Barry

## Abstract

The hippocampus has long been observed to encode a representation of an animal’s position in space. Recent evidence suggests that the nature of this representation is somewhat predictive and can be modelled by learning a successor representation (SR) between distinct positions in an environment. However, this discretisation of space is subjective making it difficult to formulate predictions about how some environmental manipulations should impact the hippocampal representation. Here we present a model of place and grid cell firing as a consequence of learning a SR from a basis set of known neurobiological features – boundary vector cells (BVCs). The model describes place cell firing as the successor features of the SR, with grid cells forming a low-dimensional representation of these successor features. We show that the place and grid cells generated using the BVC-SR model provide a good account of biological data for a variety of environmental manipulations, including dimensional stretches, barrier insertions, and the influence of environmental geometry on the hippocampal representation of space.

## Introduction

The hippocampal formation plays a central role in the ability of humans and other mammals to navigate physical space (Scoville & Milner, 1957; Morris, Garrud, Rawlins, & O’Keefe, 1982). Consistent with behavioural findings, electrophysiological studies in rodents have uncovered a range of spatially modulated neurons - yielding important insights into how the brain represents space - including place cells (O’Keefe & Dostrovsky, 1971), grid cells (Hafting, Fyhn, Molden, Moser, & Moser, 2005), head direction cells (Taube, Muller, & Ranck, 1990), and boundary vector cells (BVCs) (Barry et al., 2006; Lever, Burton, Jeewajee, O’Keefe, & Burgess, 2009; Solstad, Boccara, Kropff, Moser, & Moser, 2008). Yet how these neural representations combine to facilitate flexible and efficient goal-directed navigation, such as that observed in mammals (Etienne & Jeffery, 2004), remains an open question.

One way is to approach this problem from the field of reinforcement learning. Reinforcement learning (RL) (Sutton & Barto, 2018) seeks to address how an agent should act optimally in order to maximise expected future reward. Consequently, a quantity often used in RL is the value *V* of a state *s* in the environment which is defined as the expected cumulative reward *R*, exponentially discounted into the future by a discount parameter *γ* ∈ [0,1].

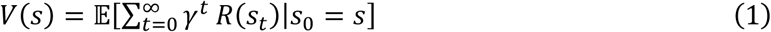

This equation can be rewritten by deconstructing value into the long-run transition statistics and corresponding reward statistics of the environment (Dayan, 1993). Here the transition statistics, denoted by *M*, is called the successor representation (SR) which represents the discounted expected future occupancy of each state *s*^′^ from the current state *s*.

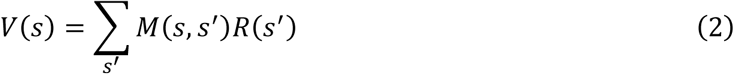

The successor representation *M* encapsulates both the short and long-term state-transition dynamics of the environment, with a time-horizon dictated by the discount parameter *γ*. Furthermore, changes to the transition and reward structure can be incorporated into the value estimates *V*(*s*) by adjusting *M* and *R* respectively. These adjustments can be made experientially using a temporal-difference learning rule, which uses the difference between predicted outcomes and the actual outcomes to improve the accuracy of the predicted estimate (O’Doherty, Dayan, Friston, Critchley, & Dolan, 2003). Thus, the SR allows the value of possible future states to be calculated flexibly and efficiently. Consequently it has been proposed that the hippocampus encodes a successor representation of space (Stachenfeld, Botvinick, & Gershman, 2017) - a claim that is further evidenced by the SR providing a good account of experimental observations of both place and grid cells. This formulation of the SR typically involves discretisation of the environment into a grid of locations, within which the SR can be learnt by transitioning around the grid of states. However, this fixed grid-world renders it hard to make predictions about how environmental manipulations, such as dimensional stretches, would immediately affect hippocampal representations. Further, in very large state spaces, estimating the SR for every state becomes an increasingly difficult and costly task. Instead, using a set of features to approximate location would allow generalisation across similar states and circumvent this curse of dimensionality. Indeed, it is clear from electrophysiological studies of the neural circuits supporting navigation that the brain does not represent space as a grid of discrete states, but rather uses an array of spatially sensitive neurons. In particular, boundary responsive neurons are found throughout the hippocampal formation, including ‘border cells’ in superficial medial entorhinal cortex (mEC) (Solstad et al., 2008) and boundary vector cells (BVCs) in subiculum (Hartley, Burgess, Lever, Cacucci, & Keefe, 2000; Barry et al., 2006; Lever et al., 2009). Since these neurons effectively provide a representation of the environmental topography surrounding the animal and – in the case of the mEC - are positioned to provide input to the main hippocampal subfields (Zhang et al., 2014), it seems plausible that they might function as an efficient substrate for a successor representation.

Thus the aim of this paper is to build and evaluate a biologically plausible successor representation based upon the firing rates of known neurobiological features in the form of boundary vector cells (BVCs) (Hartley et al., 2000; Barry et al., 2006; Solstad et al., 2008; Lever et al., 2009). Not only does this provide an efficient foundation for solving goal-directed spatial navigation problems, we show it provides an explanation for electrophysiological phenomena currently unaccounted for by the standard SR model (Stachenfeld et al., 2017).

## Model

We generate a population of BVCs following the specification used in previous iterations of the BVC model (Hartley et al., 2000; Barry & Burgess, 2007; Grieves, Duvelle, & Dudchenko, 2018). That is, the firing of the i^th^ BVC, tuned to preferred distance *d*_*i*_ and angle *ϕ*_*i*_ to a boundary at distance r and direction θ subtending at an angle *δθ* is given by:

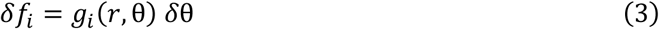

where:

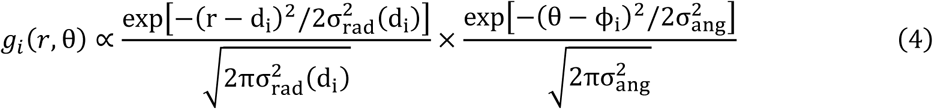

In the model, the angular tuning width σ_*ang*_ is constant and radial tuning width increases linearly with the preferred tuning distance: σ_*rad*_ (*d*_*i*_) = *d*_*i*_/β + ξ for constants β and ξ.

Using a set of *n* BVC’s, each position or state *s* in the environment corresponds to a vector of BVC firing rates ***f***(*s*) = [*f*_1_(*s*), *f*_2_(*s*), …, *f*_*n*_(*s*)] (Figure 1). We use a tilde ∼ to indicate variables constructed in the BVC feature space of ***f***. By learning a successor representation 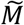 among these BVC features we can use linear function approximation of the value function to learn a set of weights 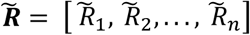 such that:

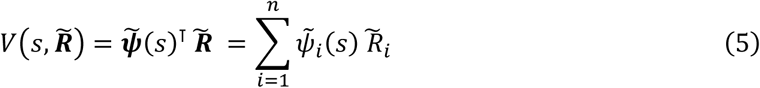

**Figure 1:**
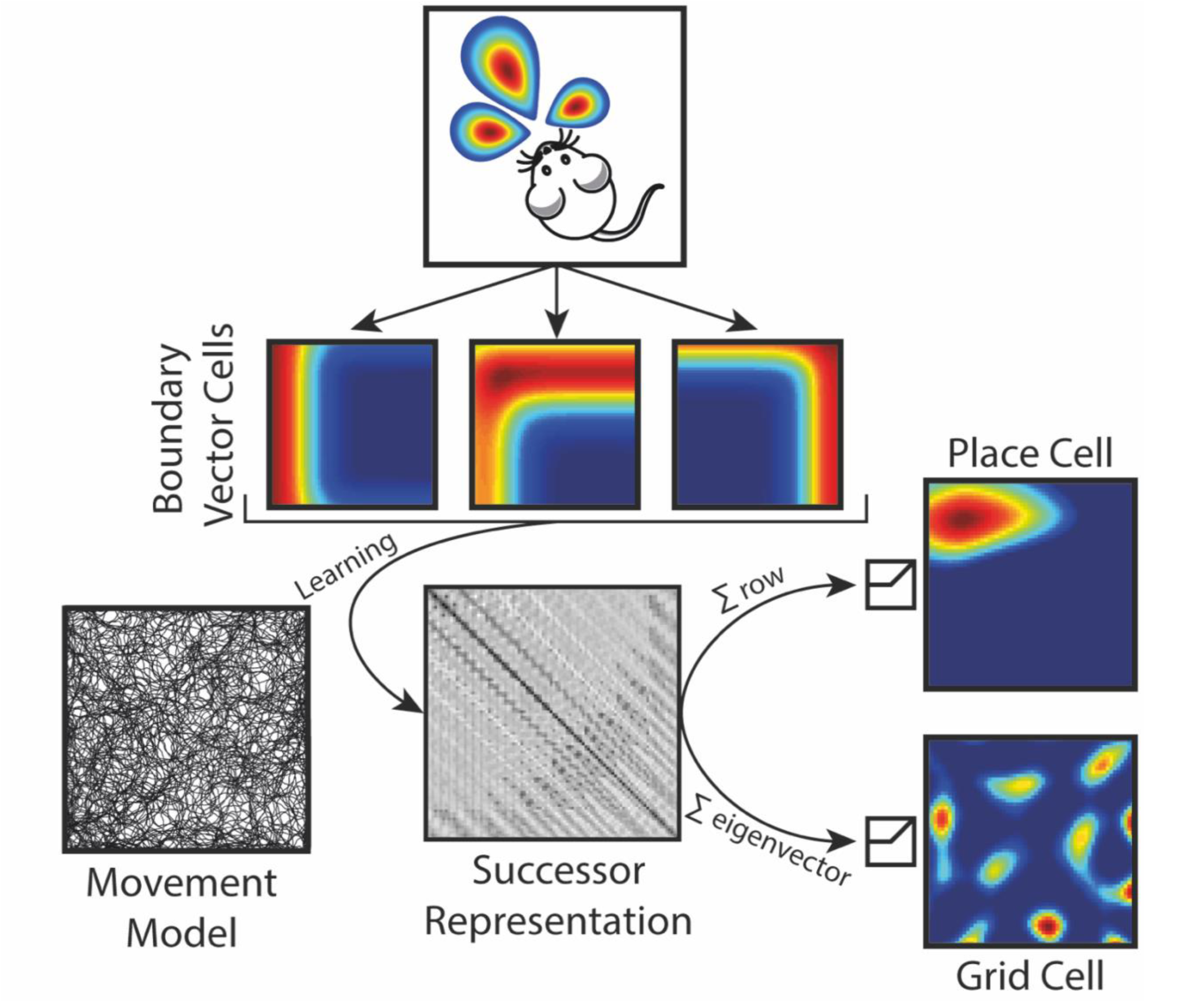
Schematic of model. Boundary vector cells (BVCs), which track the agent’s allocentric distance and direction from environmental boundaries, are used as basis features for a successor representation 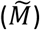. The agent’s behaviour is generated using a rodent-like movement model with the successor matrix being updated incrementally at each 50Hz time step. Following from previous analyses of the successor matrix - thresholded sums of the BVC features, weighted by rows of the SR matrix, yield unimodal firing fields with characteristics similar to CA1 place cells. Similarly, thresholded eigenvectors of the successor matrix reveal spatially periodic firing patterns similar to medial entorhinal grid cells.

Where ⊤ denotes the transpose and 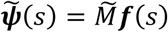 is the vector of successor features constructed using the BVCs as basis features. Analogous to the discrete state-space case where the successor matrix *M* provides a predictive mapping from the current state to the expected future states, the successor matrix 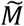 provides a predictive mapping from current BVC firing rates ***f***(*s*) to expected future BVC firing rates. Importantly, 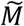 and 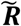 can be learnt online using temporal-difference learning rules:

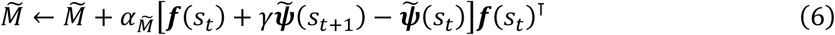

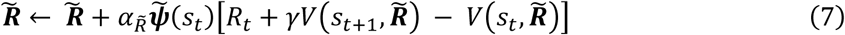

where 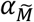 and 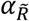 are the learning rates for the successor representation 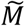 and weight vector 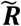 respectively. Since equation 6 is independent of reward *R*_*t*_, the model is still able to capture the structure of the environment in the absence of reward 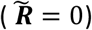 by learning the successor matrix 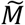. In this manner it inherently describes spatial latent learning as described in rodents (Tolman, 1948).

Consequently, we can learn through experience which BVCs are predictive of others by estimating the SR matrix 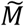. More precisely, given the agent is at position *s* with BVC population firing rate vector 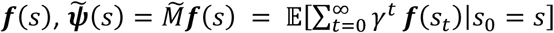 represents the expected sum of future population firing rate vectors, exponentially discounted into the future by the parameter *γ* ∈ [0,1].

This contrasts with previous implementations of the SR where rows and columns of the matrix *M* correspond to particular states. Here, rows and columns of the SR matrix correspond to particular BVCs instead. Specifically, the element 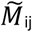 can be thought of as a weighting for how much the j^th^ BVC predicts the firing of the i^th^ BVC in the near future. Thus, whilst BVC firing ***f*** depends on the environmental boundaries, the SR matrix 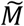 and consequently successor features 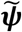 are policy dependent meaning they are shaped by behaviour. Here, in order to generate the trajectories used for learning, we utilised a motion model designed to mimic the foraging behaviour of rodents (Raudies & Hasselmo, 2012). Trajectories were sampled at a frequency of 50hz and the learning update from equation 6 was processed at every time point. All of the simulations presented here investigate the learning of successor matrix 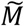 in the absence of reward 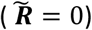.

Similar to the BVC model (Hartley et al., 2000), the firing of each simulated place cell F_i_ in a given location s is proportional to the thresholded, weighted sum of the BVCs connected to it:

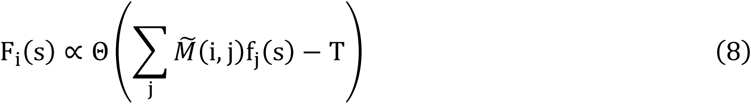

where T is the cell’s threshold and

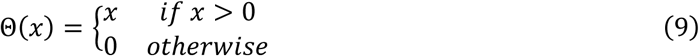

The weights in the sum (equation 8) correspond to a row of the SR matrix 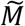 and refer to the individual contributions that a particular BVC (encoded by that row) will fire in the near future. Thus, assuming homogeneous behaviour, sets of BVCs with overlapping fields will typically exhibit mutually strong positive weights, resulting in the formation of place fields at their intersection (Figure 2a). The place cell threshold T was set to 80% of the cell’s maximum activation.

**Figure 2:**
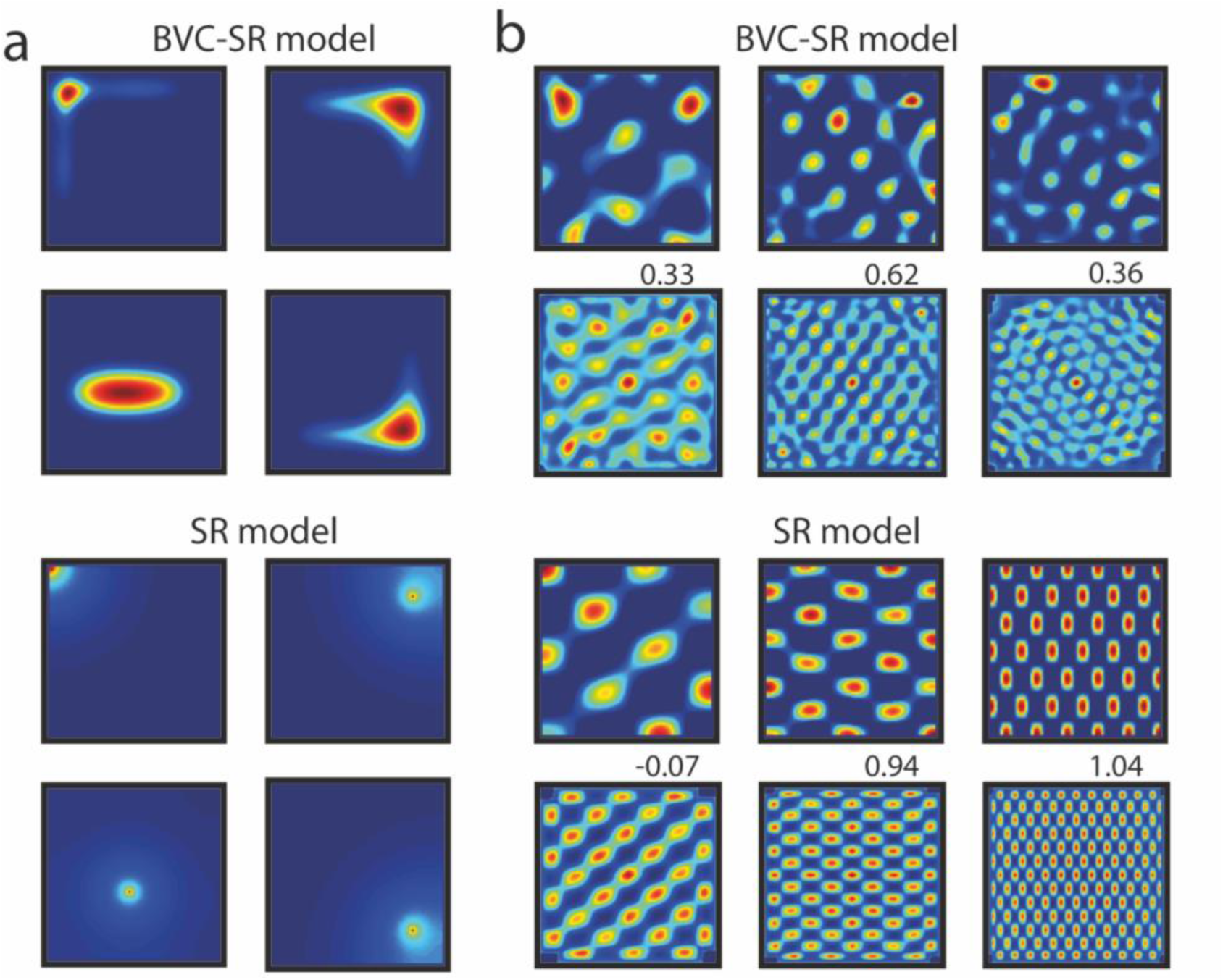
Typical place and grid cells generated by the BVC-SR and standard SR models. (**a**) Like rodent CA1 place cells, BVC-SR place cells (top) in the open field are non-uniform, irregular, and often conform to the geometry of the environment. In contrast standard SR place cells (bottom) are characterised by smooth, circular fields. (**b**) Grid cells in both the BVC-SR (1st row) and SR models (3rd row) are produced by taking the eigenvectors of the SR matrix. The corresponding spatial autocorrelograms (2^nd^ and 4^th^ rows) are used to assess the hexagonal periodicity (gridness) of the firing patterns, shown above each spatial autocorrelogram.

Grid cells in the model are generated by taking the eigendecomposition of the SR matrix 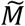 and thus represent a low-dimensional embedding of the SR. Similar to the place cells, the activity of each simulated grid cell G_i_ is proportional to a thresholded, weighted sum of BVCs. However, for the grid cells, the weights in the sum correspond to particular eigenvector 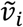 of the SR matrix 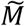 and the firing is thresholded at zero to only permit positive grid cell firing rates.

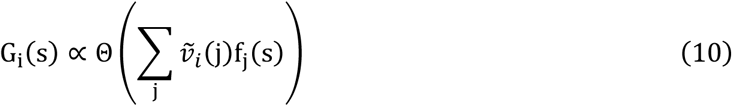

This gives rise to spatially periodic firing fields such as those observed in Figure 2b.

## Results

Following Stachenfeld and colleagues (Stachenfeld et al., 2017), we propose that the hippocampus encodes the BVC successor features 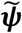 to facilitate decision making during spatial navigation. Importantly, due to the disassociation of 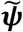 and reward weights 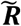 in the computation of value (equation 5), the model facilitates latent learning via the independent learning of 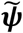 irrespective of whether reward is present. It also provides an efficient platform for goal-based navigation by simply changing the reward weights 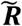.

Like real place cells and those generated by the standard SR model (Stachenfeld et al., 2017), place cells simulated with the BVC-SR model respect the transition statistics of the environment and thus do not extend through environmental boundaries. However due the nature of the underlying BVC basis features, the simulated place cells also exhibit characteristics of hippocampal place cells that are unaccounted for by the standard SR model. For example, in the standard SR model, place cell firing in a uniformly sampled open field environment tends to be characterised by circular smoothly decaying fields (Stachenfeld et al., 2017). In contrast, BVC-SR derived place fields – like real place cells and those from the BVC model (Muller, Kubie, & Ranck, 1987; Hartley et al., 2000) - are elongated along environmental boundaries and generally conform to the shape of the enclosing space (Figure 2a).

Most importantly, the use of a BVC basis set provides a means to predict how the model will respond to instantaneous changes in the structure of the environment. In Stachenfeld et al. (2017) the states available to an agent were distinct from the environmental features that constrained the allowed transitions. Thus, insertion of a barrier into an environment had no immediate effect on place or grid fields – changes in firing fields would accumulate through subsequent exploration and learning causing *M* to be updated. However, biological results indicate that place cell activity is modulated almost immediately by changes made to the geometry of an animal’s environment (O’Keefe & Burgess, 1996; Hartley et al., 2000; Barry & Burgess, 2007; Barry et al., 2006). Since BVC activity is defined relative to environmental boundaries, manipulations made to the geometry of an environment produce immediate changes in the activity of place cells without any change to the SR matrix 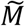. Thus, similar to the standard BVC model, elongation or compression of one or both dimensions of an open field environment distorts place cell firing in a commensurate fashion (Figure 3a,b&c), as has been seen in rodents (O’Keefe & Burgess, 1996). As a result, the basic firing properties of BVC-SR place cells – such as field size – are relatively preserved between manipulations (Figure 3d).

**Figure 3:**
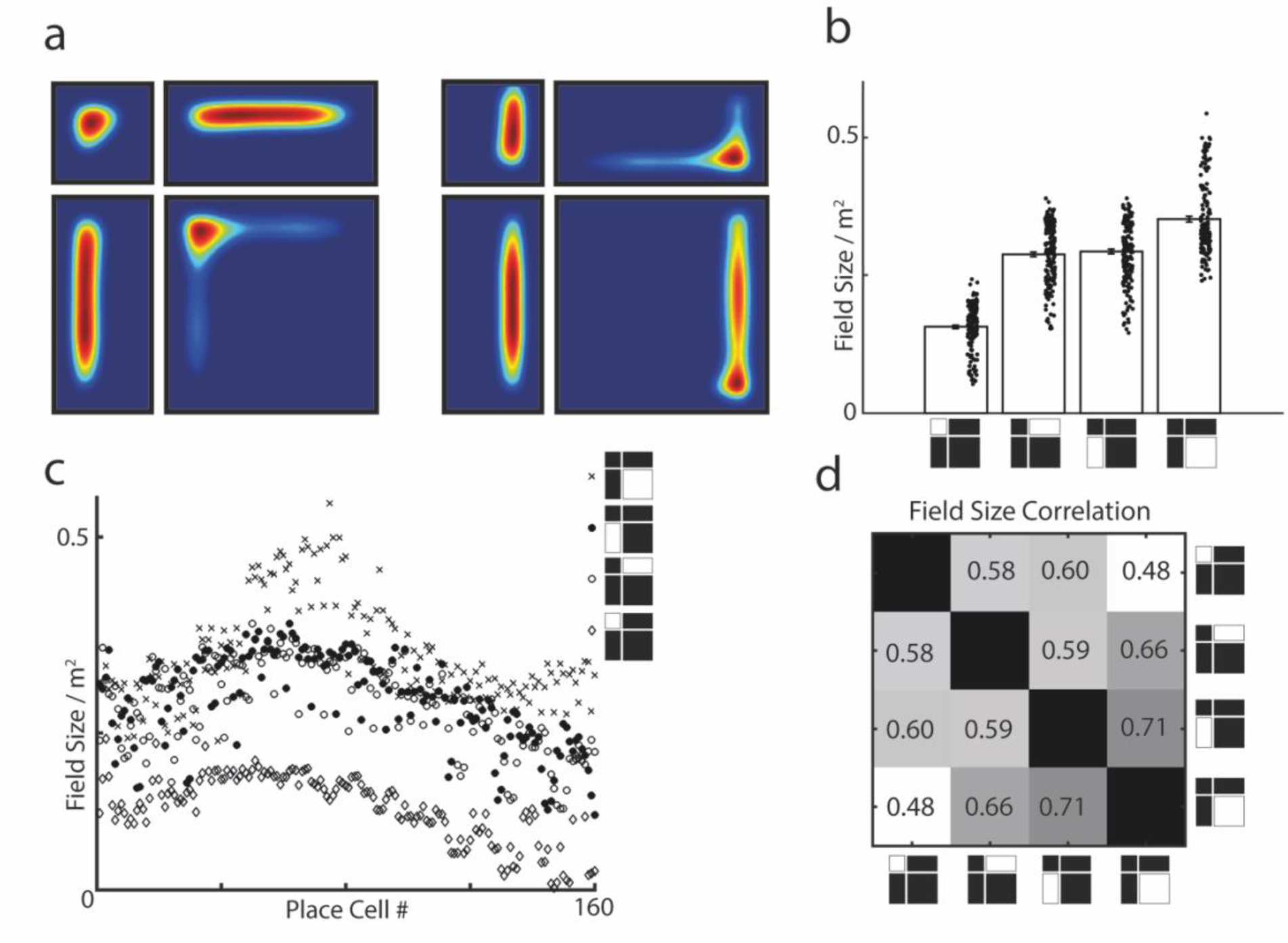
BVC-SR derived place cells deform in response to geometric manipulations made to the environment. Scaling one or both axes of an environment produces commensurate changes in the activity of BVC-SR place cells (**a**). Such that firing field size scales proportionally with environment size (**b**,**c**) while the relative size of place fields is largely preserved between environments – Pearson correlation coefficient shown (**d**).

The introduction of internal barriers into an environment provides a succinct test for geometric theories of spatial firing and has been studied in both experimental and theoretical settings. Indeed, the predictable allocentric responses of biological BVCs to inserted barriers provides some of the most compelling evidence for their existence (Lever et al., 2009; Poulter, Hartley, & Lever, 2018). In CA1 place cells, barrier insertion promotes an almost immediate duplication of place fields (Muller & Kubie, 1987) which may then be then lost or stabilised during subsequent exploration (Barry et al., 2006; Barry & Burgess, 2007). The BVC-SR model provided a good account of empirical data, exhibiting similar dynamic responses. Barrier insertion caused 23% of place cells (32 of 160) to immediately form an additional field, one being present on either side of the barrier (Figure 4a). Following further exploration, 19% of these (7 out of 32) gradually lost one of the duplicates – a modification reflecting updates made to 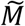 resulting from changes in behaviour due to the barrier (Figure 4b) (Barry & Burgess, 2007). Upon removal of the barrier, the simulated place cells reverted more or less to their initial tuning fields prior to barrier insertion, with minor differences due to the updated successor representation 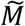.

**Figure 4:**
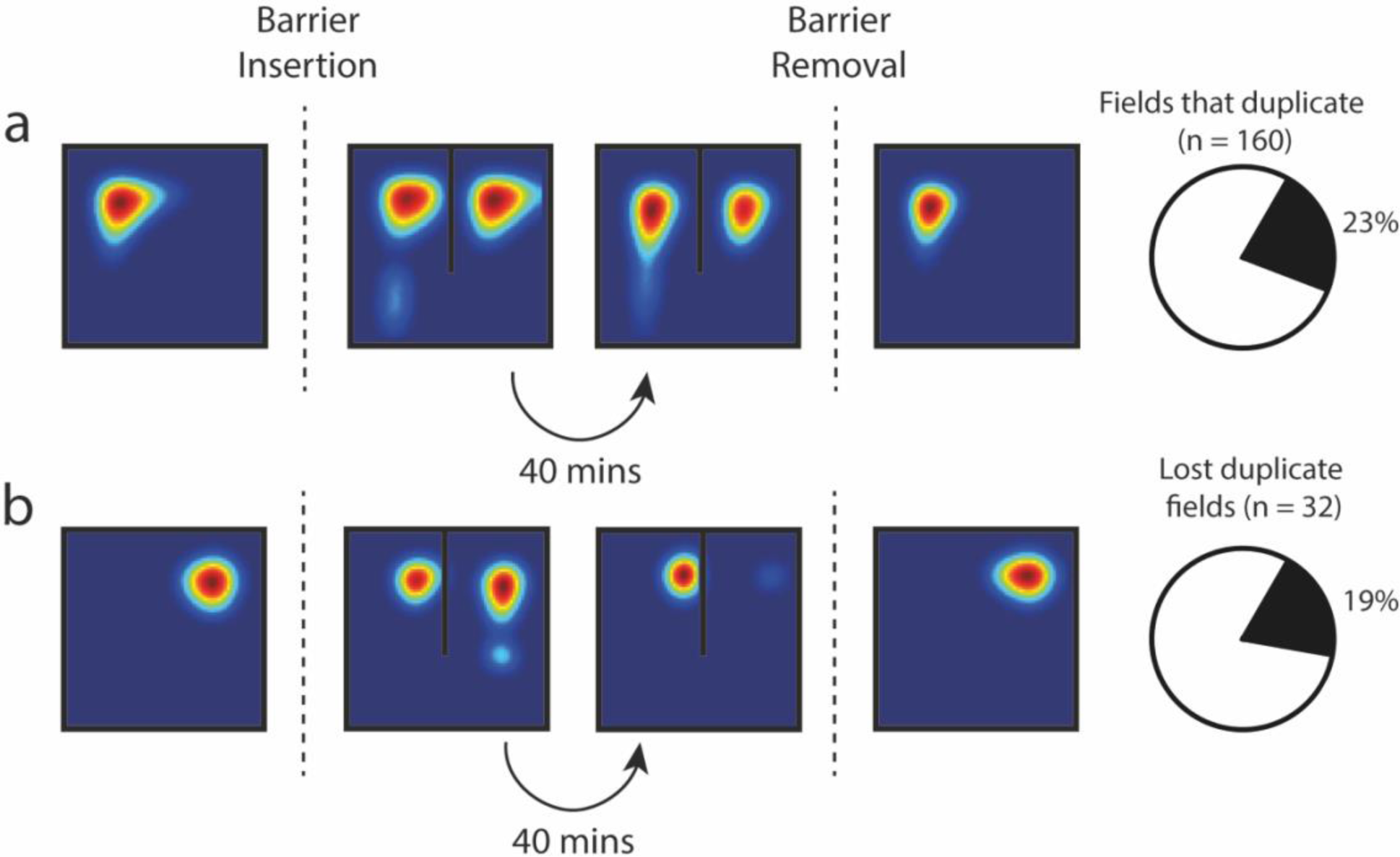
Insertion of an additional barrier into an environment can induce duplication of BVC-SR place fields. (**a**) In 23% of place cells, barrier insertion causes immediate place field duplication. In most cases (81%) the duplicate field persists for the equivalent of 40 minutes of random foraging (learning update occurs at 50Hz) (**b**) In some cases (19%) one of the duplicate fields – not necessarily the new one – is lost during subsequent exploration. Similar results have been observed *in vivo* (Barry et al., 2006).

Stachenfeld et al. (2017) previously demonstrated that eigendecomposition of the successor matrix *M* produced spatially periodic firing fields resembling mEC grid cells. Examining the eigenvectors of 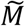, from the BVC-SR model, we found that these too resembled the regular firing patterns of grid cells (Figure 2b). Indeed, while there was no difference in the hexagonal regularity of BVC-SR and standard-SR eigenvectors (mean gridness ± SD: -0.28 ± 0.35 vs. -0.27 ± 0.60; t(318)=0.14, p=0.886), the eigenvectors from the BVC-SR exhibit less elliptic grid fields (mean field ellipticity ± SD: 0.59 ± 0.23 vs. 0.75 ± 0.25; t(318)=-5.93; p<0.001; Supplementary Figure 1) and a larger variability in field firing rates (mean coefficient of variability ± SD: 0.48 ± 0.11 vs 0.14 ± 0.11; t(318)=26.5; p<0.001; Supplementary Figure 2), similar to that observed in real grid cells (ellipticity: 0.55 ± 0.02 Krupic, Bauza, Burton, Barry, & O’Keefe, 2015; coefficient of variability: 0.58 ± 0.01 Ismakov, Barak, Jeffery, & Derdikman, 2017) – although neither yield exclusively hexagonal patterns.

Empirical work has shown that grid-patterns are modulated by environmental geometry, the regular spatial activity becoming distorted in strongly polarised environments (Derdikman et al., 2009; Krupic et al., 2015; Stensola, Stensola, Moser, & Moser, 2015). Grid-patterns derived from the standard-SR eigenvectors also exhibit distortions comparable to those seen experimentally. Thus, we next examined the regularity of BVC-SR eigenvectors derived from SR matrices trained in square and trapezoid environments. As with rodent data (Krupic et al., 2015) and the standard-SR model, we found that grid-patterns in the two halves of the square environment were considerably more regular than those derived from the trapezoid (mean correlation between spatial autocorrelograms ± SD: 0.68 ± 0.18 vs. 0.47 ± 0.15, t(318) = 10.99, p<0.001; Figure 5b). Furthermore, BVC-SR eigenvectors that exceeded a shuffled gridness threshold (see supplementary methods) – and hence were classified as grid cells – were more regular in the square than the trapezoid (mean gridness ± SD: 0.37 ± 0.17 vs. 0.10 ± 0.09; t(24) = 4.87, p <0.001; Figure 5c). In particular, as had previously been noted in rodents (Krupic et al., 2015), the regularity of these ‘grid cells’ was markedly reduced in the narrow end of the trapezoid compared to the broad end (mean gridness ± SD: -0.30 ± 0.19 vs. 0.16 ±0.23; t(22) = -5.45, p<0.001; Figure 5d), a difference that did not exist in the two halves of the square environment (mean gridness ± SD: 0.19 ± 0.25 vs. 0.22 ± 0.36; t(26) = -0.28, p = 0.78).

**Figure 5:**
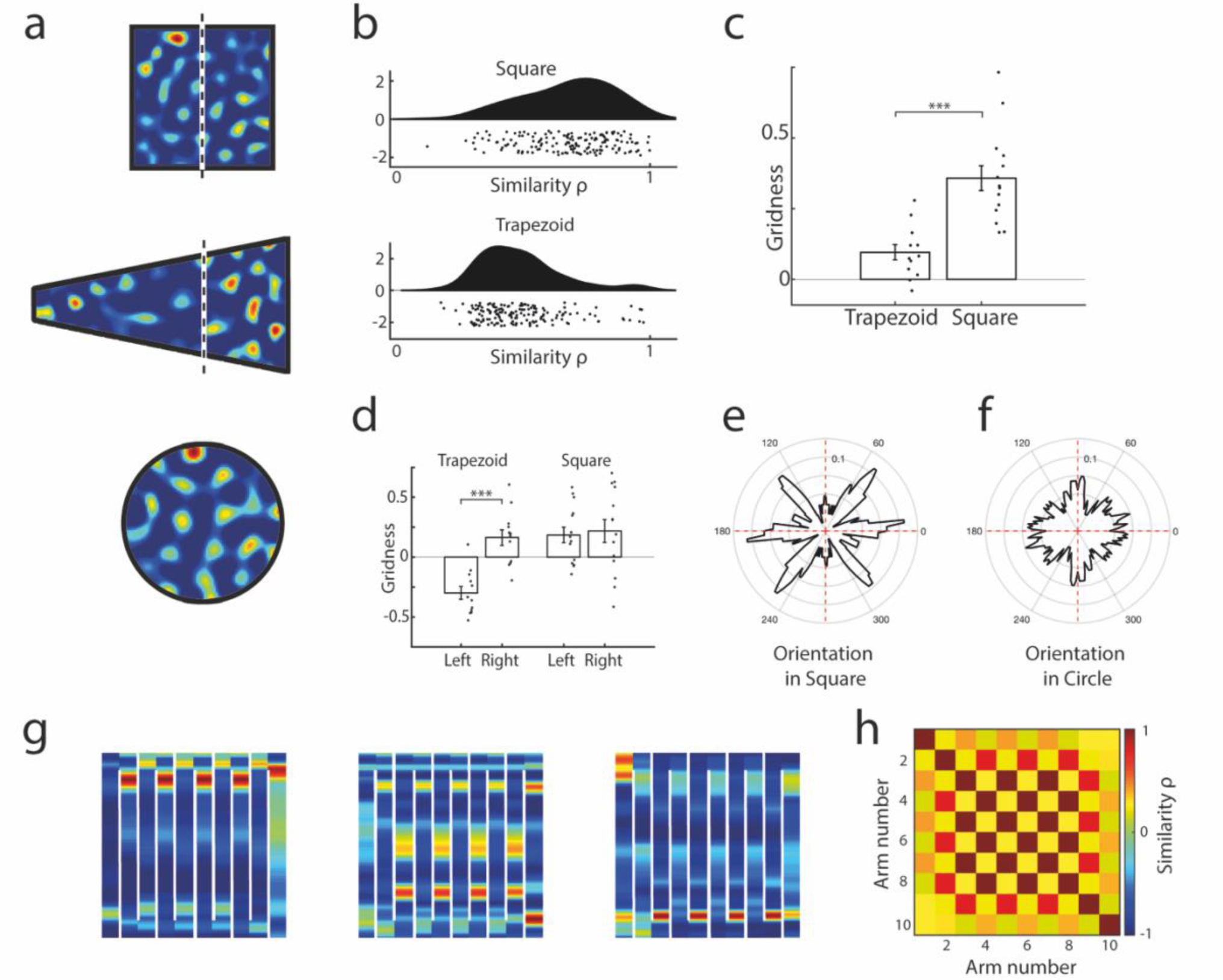
BVC-SR grid-patterns are influenced by environmental geometry. (**a**) Eigenvectors of the BVC-SR can be used to model grid cells firing patterns in a variety of different shaped enclosures (white line indicates division of square and trapezoid into halves of equal area). (**b**) Grid-patterns are more similar in the two halves of the square environment than in the two halves of the trapezoid (mean Pearson’s correlation between spatial autocorrelograms ± SD: 0.68 ± 0.18 vs. 0.47 ± 0.15, t(317) = 10.99, p<0.001), similar results have been noted in rodents (Krupic et al., 2015). (**c**) ‘Grid cells’ (grid-patterns that exceed a shuffled gridness criteria, see supplementary methods) are more hexagonal in the square environment than the trapezoid (mean gridness ± SD: 0.37 ± 0.17 vs. 0.10 ± 0.09; t(24) = 4.87, p <0.001), (**d**) the narrow half of the trapezoid being less regular than the wider end (mean gridness ± SD: -0.30 ± 0.19 vs. 0.16 ±0.23; t(22) = -5.45, p<0.001). The axes of ‘grid cells’ are more polarised (less uniform) in a square (**e**) than circular environment (**f**) (D_KL_(Square||Uniform) =0.17, D_KL_(Circle||Uniform) = 0.04; Bayes Factor = 1.00×10^−6^). (**g**) The BVC-SR eigenvector grid patterns are fragmented in a compartmentalised maze and repeat across alternating maze arms as has been observed in rodents (Derdikman et al., 2009). (**h**) The Pearson’s correlation matrix between the grid patterns on different arms of the maze has a checkerboard-like appearance due to the strong similarity between alternating internal channels of the maze (n=160 eigenvectors). Again, similar results have been noted empirically (Derdikman et al., 2009).

Rodent grid-patterns have been shown to orient relative to straight environmental boundaries – tending to align to the walls of square but not circular environments (Krupic et al., 2015; Stensola et al., 2015). In a similar vein, we saw that firing patterns of simulated grid cells also were more polarised in a square than a circular environment, tending to cluster around specific orientations (Figure 5ef). To illustrate this, we used the Kullback-Leibler divergence (D_KL_) to measure the difference between the distribution of grid orientations and a uniform distribution (see supplementary methods). We found the grid orientations in the circular environment were much closer to uniform (D_KL_(Circle||Uniform) = 0.04 vs. D_KL_(Square||Uniform) =0.17), and significantly better explained by an underlying uniform distribution as opposed to the grid orientations in the square environment (Bayes Factor = 1.00×10^6^).

Finally, the activity of grid cells recorded whilst a rodent explores a compartmentalised maze have been shown to fragment into repeated submaps for similar compartments traversed in the same direction (Derdikman et al., 2009). We examined the BVC-SR eigenvector patterns in a similar maze and found that they too fragmented into repeated submaps for alternating internal arms of the maze (Figure 5g). Consequently, the Pearson’s correlation matrix between eigenvector patterns on different arms of the maze exhibits a strong checkboard-like appearance (Figure 5h), exemplifying the repetition of alternated submaps in a manner more similar to the rodent data (Derdikman et al., 2009) than previous implementations of the SR (Stachenfeld et al., 2017).

## Discussion

The model presented here links the BVC model of place cell firing with a SR to provide an efficient platform for using reinforcement learning to navigate space. The work builds upon previous implementations of the SR by replacing the underlaying grid of states with the firing rates of known neurobiological features - BVCs, which have been observed in the hippocampal formation (Barry et al., 2006; Solstad et al., 2008; Lever et al., 2009) and can be derived from optic flow (Raudies & Hasselmo, 2012). As a consequence, the place cells generated using the BVC-SR approach presented here produce more realistic fields that conform to the shape of the environment. Unlike previous SR implementations, the BVC-SR place fields respond immediately to environmental manipulations such as dimensional stretches and barrier insertions in a similar manner to real place cells.

Comparable to previous SR implementations, the eigenvectors of the SR matrix 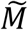 display grid cell like periodicity when projected back onto the BVC state space, with reduced periodicity in polarised enclosures such as trapezoids. Further, likely due to the experiential learning and the natural smoothness of the BVC basis features, the eigenvectors from the BVC-SR model exhibit more realistic variations among grid fields, resulting in a model of grid cells that is more similar to biological recordings than previous implementations of the SR. This form of eigendecomposition is similar to other dimensionality reduction techniques that have been used to generate grid cells from populations of idealised place cells with a generalised Hebbian learning rule (Oja, 1982; Dordek, Soudry, Meir, & Derdikman, 2016). Previously, low dimensional encodings such as these have been shown to accelerate learning and facilitate vector-based navigation (Gustafson & Daw, 2011; Banino et al., 2018).

The model extends upon the BVC model of place cell firing (Hartley et al., 2000; Barry et al., 2006; Barry & Burgess, 2007) by also providing a means of predicting how environmental boundaries might affect the firing of grid cells. Furthermore, whilst both models produce similar place cells if the agent samples the environment uniformly, the policy dependence of the BVC-SR model provides a mechanism for estimating how behavioural biases will influence place cell firing. These models both use BVCs as the basis for allocentric place representations in the brain. As a consequence, they would be unable to distinguish between visually identical compartments based on boundary information alone. To achieve this, the models would require some form of additional information about the agent’s past trajectory, such as a path integration signal. Theoretical evidence (Byrne, Becker, & Burgess, 2007; Bicanski & Burgess, 2018) suggests that recently discovered egocentric BVCs (Hinman, Chapman, & Hasselmo, 2019; Gofman et al., 2019) could provide the link between the egocentric perception of the environment to an allocentric representation in the hippocampal formation.

The focus of this work has centred on the representation of successor features in the hippocampus during the absence of environmental reward. However a key feature of SR models is their ability to adapt flexibly and efficiently to changes in the reward structure of the environment (Dayan, 1993; Russek, Momennejad, Botvinick, Gershman, & Daw, 2017; Stachenfeld et al., 2017). This is permitted by the independent updating of reward weights (equation 7) combined with its immediate effect on the computation of value (equation 5). Reward signals analogous to that used in the model have been shown to exist in the orbitofrontal cortex of rodents (Sul, Kim, Huh, Lee, & Jung, 2010), humans (Gottfried, O’Doherty, & Dolan, 2003; Kringelbach, 2005) and non-human primates (Tremblay & Schultz, 1999). Meanwhile a candidate area for integrating orbitofrontal reward representations with hippocampal successor features to compute value could be anterior cingulate cortex (Shenhav, Botvinick, & Cohen, 2013; Kolling et al., 2016). Finally, the model relies on a prediction error signal for learning both the reward weights and successor features (equations 6-7). Whilst midbrain dopamine neurons have long been considered a source for such a reward prediction error (Schultz, Dayan, & Montague, 1997), mounting evidence suggests they may also provide the sensory prediction error signal necessary for computing successor features with temporal-difference learning (Chang, Gardner, Di Tillio, & Schoenbaum, 2017; Gardner, Schoenbaum, & Gershman, 2018).

Successor features have been used to accelerate learning in tasks where transfer of knowledge is useful, such as virtual and real world navigation tasks (Barreto et al., 2017; Zhang, Springenberg, Boedecker, & Burgard, 2017). Whilst the successor features used in this paper were built upon known neurobiological spatial neurons, BVCs, the framework itself could be applied to any basis of sensory neurons that are predictive of reward in a task. Thus, the framework could be adapted to use basis features that are receptive to the frequency of auditory cues (Aronov, Nevers, & Tank, 2017), or even the size and shape of birds (Constantinescu, O’Reilly, & Behrens, 2016).

In summary, the model describes the formation of place and grid fields in terms the geometric properties and transition statistics of the environment, whilst providing an efficient platform for goal-directed spatial navigation. This has particular relevance for the neural underpinnings of spatial navigation, although the framework itself could be applied to other basis sets of sensory features.

## Supplementary Methods

All simulations were implemented in MATLAB 2018b using the same set of 160 BVCs with parameters σ_*ang*_ = 11.25°, β = 12, ξ = 8. We used 16 preferred angles (0°, 22.5°, 45°, 67.5°, 90°, 112.5°, 135°, 157.5°, 180°, 202.5°, 225°, 247.5°, 270°, 292.5°, 315°, 337.5°) at 10 preferred distances (3.3cm, 10.2cm, 17.5cm, 25.3cm, 33.7cm, 42.6cm, 52.2cm, 62.4cm, 73.3cm, 85.0cm) chosen to provide uniform overlap between consecutive angular and radial tunings. Environments were discretised into 1×1cm bins for analysis, with rate maps filtered by a 13×13 Gaussian smoothing kernel with σ = 3cm. All simulations learned the SR 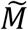 by implementing equation 8 at a rate of 50Hz with 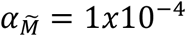 and γ = 0.995. Unless otherwise stated (such as in Figure 4), learning of 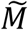 was implemented using 2 hours of random foraging trajectory simulated using a model that mimics rodent foraging (Raudies & Hasselmo, 2012).

The environment dimensions used for the simulations in Figure 3 were 60×60cm, 60×120cm, 120×60cm and 120×120cm. The simulations for Figure 4 used a 65×65cm box with the insertion of a 40cm barrier. The environments used in Figure 5 consisted of a 95×95cm box, an 80cm diameter circle, and a trapezoid formed of base lengths of 20cm and 90cm with height 172cm.

Gridness scores were calculated following (Hafting et al., 2005), with gridness thresholds used to identify the eigenvectors for analysis in Figures 5c-f identified using the 95th percentile of a field shuffling procedure (C. Barry & Burgess, 2017). The orientations of the main 3 axes of a ‘grid cell’ were identified using the central part of the spatial autocorrelogram encompassing the central peak and the 6 closest surrounding peaks. The orientations were calculated as the angles between the horizontal central axis and the 3 lines connecting the 3 closest peaks to the central peak in an anticlockwise order. Due to the symmetry in spatial autocorrelograms, the other 3 peaks have the same orientation modulo 180°. To assess the orientation clustering of grid-cells in the square and circular environments in Figure 5ef, we calculated the Kullback-Leibler (KL) divergence between the distributions of the grid orientations and a uniform distribution using a bin width of 12°.

The KL divergence *D*_*KL*_ is a measure of how different a probability distribution *P* (grid orientations) is from a reference distribution *Q* (uniform distribution) defined on the same probability space.

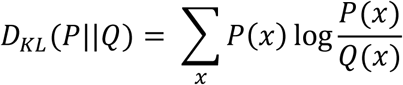

*D*_*KL*_ ≥ 0 and *D*_*KL*_ = 0 if and only if *P* = *Q*.

The Bayes Factor reported compares the likelihood of the grid orientations in the circle as a result of two models – one model being that they share the same the distribution of orientations as in the square environment, and another being that they are sampled from a uniform distribution of orientations.

For comparison with the standard SR model, the environment was discretised into 1×1cm bins and the SR matrix *M* was computed as the discounted sum of one-step transition matrices *T* describing a uniform random walk on this state-space: 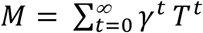 (Stachenfeld et al., 2017). As with the BVC-SR model, a discount factor of γ = 0.995 was used. The ellipticity and firing rate variability analyses were computed for the 95×95cm square environment.

The grid cell rate map ellipticity reported for the biological data (Krupic, Bauza, Burton, Barry, & O’Keefe, 2015) was calculated by fitting an ellipse to the 6 peaks surrounding the central peak of the spatial autocorrelogram. The eccentricity *e* of this ellipse used as the measure of the grid ellipticity:

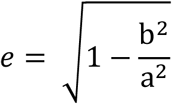

where a and b are the lengths of the longer and shorter axis of the fitted ellipse. Since neither the BVC-SR or standard SR models yield exclusively hexagonal grid patterns, ellipticity of the eigenvector rate maps was calculated by thresholding the spatial autocorrelogram at a value of 0.2 and identifying the central peak. Then the eccentricity *e* of this central peak was used as the measure of the grid ellipticity, where a and b are the lengths of the longer and shorter axis of the central peak.

Firing rate variability of the eigenvector rate maps was analysed following the method of Ismakov et al., (2017). Grid fields were identified using the watershed transform of each eigenvector rate map, and the coefficient of variability (CV) was calculated as the standard deviation of these peaks, divided by the mean of the peaks.

**Supplementary Figure 1:**
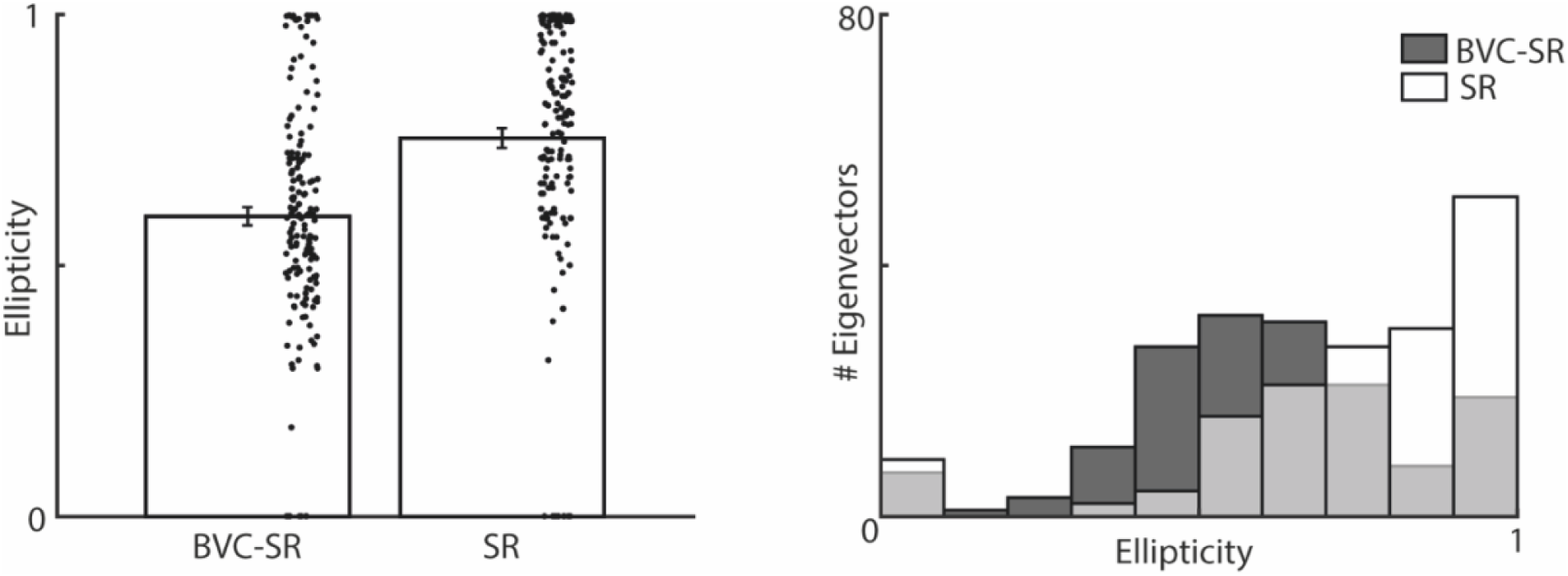
Grid fields generated using eigenvectors from the BVC-SR model are less elliptic than those from the standard SR model. Lower values indicate more circular fields and larger values indicate more elliptic fields, with a value of 0 indicating a perfect circle. a) Grid fields generated using the BVC-SR model had significantly lower ellipticity than the standard SR model (mean field ellipticity ± SD: 0.59 ± 0.23 vs. 0.75 ± 0.25; t(318)=-5.93; p<0.001), and were similar to observations of real grid cells (Krupic et al., 2015). b) Histogram of the grid field ellipticity (N=160 eigenvectors)

**Supplementary Figure 2:**
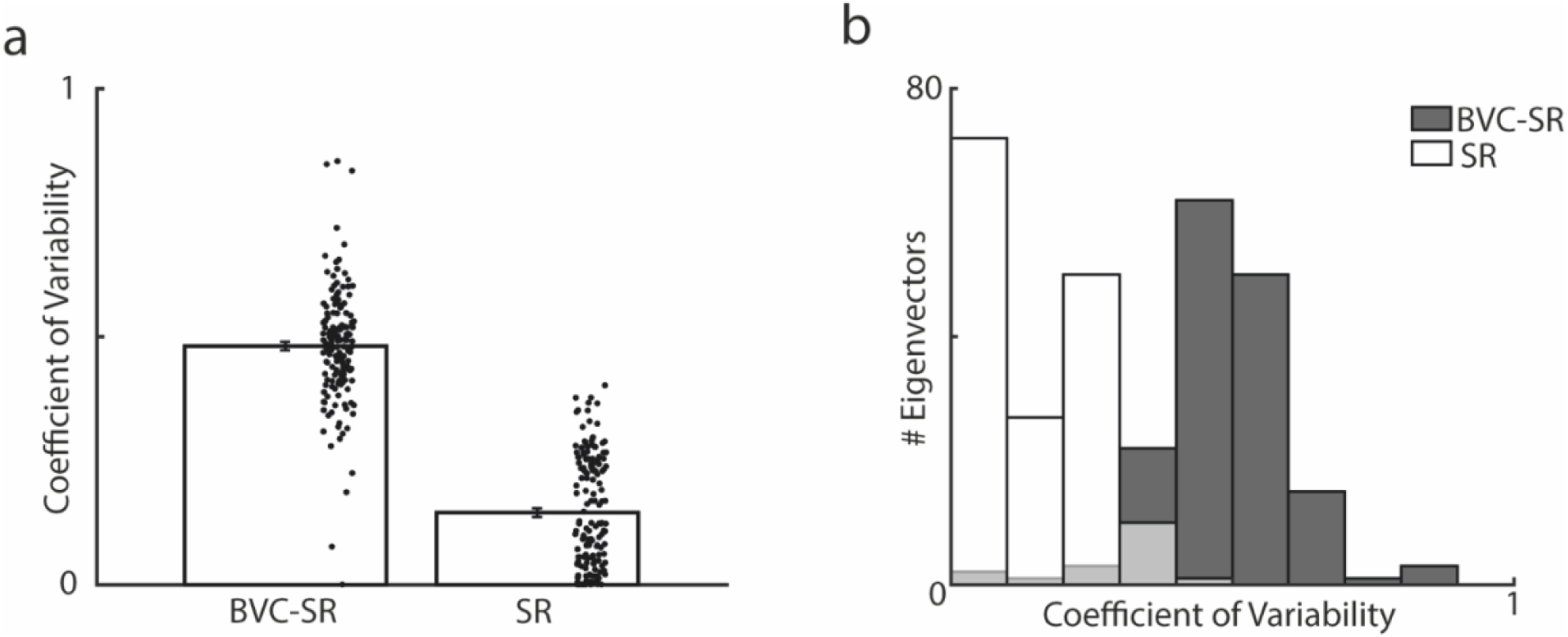
Grid fields generated using eigenvectors from the BVC-SR model exhibit more firing rate variability than the standard SR model. Following the method of Ismakov et al., (2017), the peak firing rates of grid fields was used to compute a coefficient of variability for each eigenvector (CV; SD divided by mean). a) The CV for eigenvectors produced by the BVC-SR model were significantly larger than that observed in the standard SR model (mean CV ± SD: 0.48 ± 0.11 vs 0.14 ± 0.11; t(318)=26.5; p<0.001), and similar to that observed in real grid cells (Ismakov et al., 2017). b) Histogram of the CV for each of the models (N=160 eigenvectors).

## Notes

### Competing Interest Statement

The authors have declared no competing interest.

## References

Aronov, D., Nevers, R., & Tank, D. W. (2017). Mapping of a non-spatial dimension by the hippocampal–entorhinal circuit. Nature, 543(7647), 719–722. https://doi.org/10.1038/nature21692

Banino, A., Barry, C., Uria, B., Blundell, C., Lillicrap, T., Mirowski, P., … Kumaran, D. (2018). Vector-based navigation using grid-like representations in artificial agents. Nature, 557(7705), 429–433. https://doi.org/10.1038/s41586-018-0102-6

Barreto, A., Dabney, W., Munos, R., Hunt, J. J., Schaul, T., Van Hasselt, H., & Silver, D. (2017). Successor features for transfer in reinforcement learning. Advances in Neural Information Processing Systems, *2017*-*Decem*, 4056–4066. Retrieved from https://arxiv.org/pdf/1606.05312.pdf

Barry, C., & Burgess, N. (2017). To be a Grid Cell: Shuffling procedures for determining “Gridness.” BioRxiv, 230250. https://doi.org/10.1101/230250

Barry, Caswell, & Burgess, N. (2007). Learning in a Geometric Model of Place Cell Firing. Hippocampus, 17, 786–800. https://doi.org/10.1002/hipo

Barry, Caswell, Lever, C., Hayman, R., Hartley, T., Burton, S., O’Keefe, J., … Burgess, N. (2006). The Boundary Vector Cell Model of Place Cell Firing and Spatial Memory. Reviews in the Neurosciences, 17 (1–2). https://doi.org/10.1515/REVNEURO.2006.17.1-2.71

Bicanski, A., & Burgess, N. (2018). A neural-level model of spatial memory and imagery. ELife, 7(7052), 1–3. https://doi.org/10.7554/eLife.33752

Byrne, P., Becker, S., & Burgess, N. (2007). Remembering the past and imagining the future: A neural model of spatial memory and imagery. Psychological Review, 114(2), 340–375. https://doi.org/10.1037/0033-295X.114.2.340

Chang, C. Y., Gardner, M., Di Tillio, M. G., & Schoenbaum, G. (2017). Optogenetic Blockade of Dopamine Transients Prevents Learning Induced by Changes in Reward Features. Current Biology, 27(22), 3480-3486.e3. https://doi.org/10.1016/j.cub.2017.09.049

Constantinescu, A. O., O’Reilly, J. X., & Behrens, T. E. J. (2016). Organizing conceptual knowledge in humans with a gridlike code. Science, 352(6292), 1464–1468. https://doi.org/10.1126/science.aaf0941

Dayan, P. (1993). Improving generalization for temporal difference learning: The successor representation. Neural Computation, 5(4), 613–624. https://doi.org/10.1162/neco.1993.5.4.613

Derdikman, D., Whitlock, J. R., Tsao, A., Fyhn, M., Hafting, T., Moser, M. B., & Moser, E. I. (2009). Fragmentation of grid cell maps in a multicompartment environment. Nature Neuroscience, 12(10), 1325–1332. https://doi.org/10.1038/nn.2396

Dordek, Y., Soudry, D., Meir, R., & Derdikman, D. (2016). Extracting grid cell characteristics from place cell inputs using non-negative principal component analysis. ELife, 5 (MARCH2016), 1–36. https://doi.org/10.7554/eLife.10094

Etienne, A. S., & Jeffery, K. J. (2004). Path integration in mammals. Hippocampus, Vol. 14, pp. 180–192. https://doi.org/10.1002/hipo.10173

Gardner, M. P. H., Schoenbaum, G., & Gershman, S. J. (2018). Rethinking dopamine as generalized prediction error. Proceedings of the Royal Society B: Biological Sciences, 285(1891). https://doi.org/10.1098/rspb.2018.1645

Gofman, X., Tocker, G., Weiss, S., Boccara, C. N., Lu, L., Moser, M. B., … Derdikman, D. (2019). Dissociation between Postrhinal Cortex and Downstream Parahippocampal Regions in the Representation of Egocentric Boundaries. Current Biology, 29(16), 2751-2757.e4. https://doi.org/10.1016/j.cub.2019.07.007

Gottfried, J. A., O’Doherty, J., & Dolan, R. J. (2003). Encoding predictive reward value in human amygdala and orbitofrontal cortex. Science, 301(5636), 1104–1107. https://doi.org/10.1126/science.1087919

Grieves, R. M., Duvelle, É., & Dudchenko, P. A. (2018). A boundary vector cell model of place field repetition. Spatial Cognition and Computation, 18(3), 217–256. https://doi.org/10.1080/13875868.2018.1437621

Gustafson, N. J., & Daw, N. D. (2011). Grid cells, place cells, and geodesic generalization for spatial reinforcement learning. PLoS Computational Biology, 7(10). https://doi.org/10.1371/journal.pcbi.1002235

Hafting, T., Fyhn, M., Molden, S., Moser, M.-B., & Moser, E. I. (2005). Microstructure of a spatial map in the entorhinal cortex. Nature, 436(7052), 801–806. https://doi.org/10.1038/nature03721

Hartley, T., Burgess, N., Lever, C., Cacucci, F., & Keefe, J. O. (2000). Modeling Place Fields in Terms of the Cortical Inputs to the Hippocampus. Hippocampus, 379, 369–379.

Hinman, J. R., Chapman, G. W., & Hasselmo, M. E. (2019). Neuronal representation of environmental boundaries in egocentric coordinates. Nature Communications, 10(1), 1–8. https://doi.org/10.1038/s41467-019-10722-y

Ismakov, R., Barak, O., Jeffery, K., & Derdikman, D. (2017). Grid Cells Encode Local Positional Information. Current Biology, 27(15), 2337-2343.e3. https://doi.org/10.1016/j.cub.2017.06.034

Kolling, N., Wittmann, M. K., Behrens, T. E. J., Boorman, E. D., Mars, R. B., & Rushworth, M. F. S. (2016, October 1). Value, search, persistence and model updating in anterior cingulate cortex. Nature Neuroscience, Vol. 19, pp. 1280–1285. https://doi.org/10.1038/nn.4382

Kringelbach, M. L. (2005, September). The human orbitofrontal cortex: Linking reward to hedonic experience. Nature Reviews Neuroscience, Vol. 6, pp. 691–702. https://doi.org/10.1038/nrn1747

Krupic, J., Bauza, M., Burton, S., Barry, C., & O’Keefe, J. (2015). Grid cell symmetry is shaped by environmental geometry. Nature, 518(7538), 232–235. https://doi.org/10.1038/nature14153

Lever, C., Burton, S., Jeewajee, A., O’Keefe, J., & Burgess, N. (2009). Boundary Vector Cells in the Subiculum of the Hippocampal Formation. Journal of Neuroscience, 29(31), 9771–9777. https://doi.org/10.1523/JNEUROSCI.1319-09.2009

Morris, R. G. M., Garrud, P., Rawlins, J. N. P., & O’Keefe, J. (1982). Place navigation impaired in rats with hippocampal lesions. Nature, 297(5868), 681–683. https://doi.org/10.1038/297681a0

Muller, R., Kubie, J., & Ranck, J. (1987). Spatial firing patterns of hippocampal complex-spike cells in a fixed environment. The Journal of Neuroscience, 7(7), 1935–1950. https://doi.org/10.1523/JNEUROSCI.07-07-01935.1987

Muller, R. U., & Kubie, J. L. (1987). The Effects of Changes in the Environment Hippocampal Cells on the Spatial Firing of. The Journal of Neuroscience, 7(July).

O’Doherty, J. P., Dayan, P., Friston, K., Critchley, H., & Dolan, R. J. (2003). Temporal difference models and reward-related learning in the human brain. Neuron, 38(2), 329–337. https://doi.org/10.1016/S0896-6273(03)00169-7

O’Keefe, J., & Burgess, N. (1996). Geometric determinants of the place fields of hippocampal neurons. Nature, Vol. 381, pp. 425–428. https://doi.org/10.1038/381425a0

O’Keefe, J., & Dostrovsky, J. (1971). The hippocampus as a spatial map. Preliminary evidence from unit activity in the freely-moving rat. Brain Research, 34(1), 171–175.

Oja, E. (1982). A Simplified Neuron Model as a Principal Component Analyzer. In Journal of Mathematical Biology (Vol. 15). Springer-Verlag.

Poulter, S., Hartley, T., & Lever, C. (2018). The Neurobiology of Mammalian Navigation. Current Biology, 28(17), R1023–R1042. https://doi.org/10.1016/j.cub.2018.05.050

Raudies, F., & Hasselmo, M. E. (2012). Modeling Boundary Vector Cell Firing Given Optic Flow as a Cue. PLoS Comput Biol, 8(6), 1002553. https://doi.org/10.1371/journal.pcbi.1002553

Russek, E. M., Momennejad, I., Botvinick, M. M., Gershman, S. J., & Daw, N. D. (2017). Predictive representations can link model-based reinforcement learning to model-free mechanisms. PLoS Computational Biology, 13(9). https://doi.org/10.1371/journal.pcbi.1005768

Schultz, W., Dayan, P., & Montague, P. R. (1997). A Neural Substrate of Prediction and Reward. Science, 275(5306), 1593–1599. https://doi.org/10.1126/science.275.5306.1593

Scoville, W. B., & Milner, B. (1957). Loss of recent memory after bilateral hippocampal lesions. Journal of Neurology, Neurosurgery, and Psychiatry, 20(1), 11–21. https://doi.org/10.1136/jnnp.20.1.11

Shenhav, A., Botvinick, M. M., & Cohen, J. D. (2013, July 24). The expected value of control: An integrative theory of anterior cingulate cortex function. Neuron, Vol. 79, pp. 217–240. https://doi.org/10.1016/j.neuron.2013.07.007

Solstad, T., Boccara, C. N., Kropff, E., Moser, M. B., & Moser, E. I. (2008). Representation of geometric borders in the entorhinal cortex. Science, 322(5909), 1865–1868. https://doi.org/10.1126/science.1166466

Stachenfeld, K. L., Botvinick, M. M., & Gershman, S. J. (2017). The hippocampus as a predictive map. Nature Neuroscience. https://doi.org/10.1038/nn.4650

Stensola, T., Stensola, H., Moser, M. B., & Moser, E. I. (2015). Shearing-induced asymmetry in entorhinal grid cells. Nature, 518(7538), 207–212. https://doi.org/10.1038/nature14151

Sul, J. H., Kim, H., Huh, N., Lee, D., & Jung, M. W. (2010). Distinct roles of rodent orbitofrontal and medial prefrontal cortex in decision making. Neuron, 66(3), 449–460. https://doi.org/10.1016/j.neuron.2010.03.033

Sutton, R. S., & Barto, A. G. (2018). Reinforcement Learning: An Introduction (2nd ed.). MIT Press.

Taube, J. S., Muller, R. U., & Ranck, J. B. (1990). Head-direction cells recorded from the postsubiculum in freely moving rats. I. Description and quantitative analysis. The Journal of Neuroscience : The Official Journal of the Society for Neuroscience, 10(2), 420–435. https://doi.org/10.1212/01.wnl.0000299117.48935.2e

Tolman, E. C. (1948). Cognitive maps in rats and men. Psychological Review, Vol. 55, pp. 189–208. https://doi.org/10.1037/h0061626

Tremblay, L., & Schultz, W. (1999). Relative reward preference in primate orbitofrontal cortex. Nature, 398(6729), 704–708. https://doi.org/10.1038/19525

Zhang, J., Springenberg, J. T., Boedecker, J., & Burgard, W. (2017). Deep reinforcement learning with successor features for navigation across similar environments. IEEE International Conference on Intelligent Robots and Systems, *2017*-*Septe*, 2371–2378. https://doi.org/10.1109/IROS.2017.8206049

Zhang, S. J., Ye, J., Couey, J. J., Witter, M., Moser, E. I., & Moser, M. B. (2014, February 5). Functional connectivity of the entorhinal - Hippocampal space circuit. Philosophical Transactions of the Royal Society B: Biological Sciences, Vol. 369. https://doi.org/10.1098/rstb.2012.0516

